# Differential expression of transposable elements in stem cell lineages of the preimplantation embryo

**DOI:** 10.1101/2020.09.13.295501

**Authors:** Eddie Dai, Nehemiah S. Alvarez, M. A. Karim Rumi

**Affiliations:** Department of Pathology and Laboratory Medicine, University of Kansas Medical Center, Kansas City, Kansas, USA; Department of Molecular and Integrative Physiology, University of Kansas Medical Center, Kansas City, Kansas, USA; Institute for Reproduction and Perinatal Research, University of Kansas Medical Center, Kansas City, Kansas, USA

**Keywords:** Transposable element, ES cells, TS cells, XEN cells

## Abstract

Approximately half of the human genome is comprised of transposable elements (TEs), which are genetic elements capable of amplifying themselves within the genome. Throughout the course of human life, TEs are expressed in germ cells, the preimplantation embryo, and the placenta but silenced elsewhere. However, the functions of TEs during embryonic development are poorly understood. Trophoblast stem (TS), embryonic stem (ES), and extraembryonic endoderm stem (XEN) cells are cell lineages derived from the preimplantation embryo and known to have different TE silencing mechanisms. Thus, it is likely distinct TEs are expressed in each lineage and that proteins coded by these TEs have lineage-specific functions. The purpose of this research was to determine which TEs are expressed in each of these stem cell lineages and to compare expression levels between lineages. Each lineage’s transcriptome was analyzed by quantifying TE expression in RNA-sequencing data from mouse stem cells. Expression data were then used for differential expression analyses performed between the cell types. It was found that certain families of TEs are distinctly expressed in certain lineages, suggesting expression of these families may be involved in the differentiation and development of each lineage, the understanding of which can lead to improved stem cell therapies and capacity to study human embryonic development.

## 1. INTRODUCTION

Within humans, only 5% of the genome is comprised of protein-coding sequences while about half of the genome is composed of transposable elements (TEs) (Lander et al., 2001). TEs are classified as genetic elements capable of moving from place to place within a genome (Kazazian, 2004). Autonomously replicating TEs include long terminal repeat (LTR) retrotransposons, more commonly known as endogenous retroviruses (ERVs), and long interspersed nuclear elements (LINEs). Because of their replication capability, TEs exist in high copy numbers throughout the human genome (Feschotte and Gilbert, 2012; Gifford and Tristem, 2003; Jern and Coffin, 2008).

TE expression is largely repressed in somatic cells. However, germ cells, the preimplantation embryo, and the placenta are permissive for TE expression (Meyer et al., 2017). Three stem cell lineages comprise the preimplantation embryo: trophoblast stem (TS) cells, embryonic stem (ES) cells, and extraembryonic endoderm stem (XEN) cells. TS cells originate in the trophectoderm (TE) and, along with the endometrium, make the placenta, ES cells originate in the epiblast (EPI) and form the embryo proper, and XEN cells originate in the primitive endoderm (PrE) and make the yolk sac (Wamaitha and Niakan, 2018).

The study of TEs in these stem cell lineages has the potential to reveal their function in mammalian development. Evidence suggest that pluripotency regulatory networks arose due to retrotransposition events that resulted in the insertion of transcription factor (TF) binding sites upstream of transcription start sites (Jacques et al., 2013). Additionally, growing evidence has shown that TE insertions can provide the raw genetic material for the emergence of protein-coding genes which can take on essential cellular function (Joly-Lopez and Bureau, 2018; Naville et al., 2016). One such instance of this is the co-option of ERV env-proteins by mammalian ancestors which contributed to the evolution of the placenta (Blaise et al., 2005).

Identifying the TE expression patterns in early stem cell lineages will establish an important foundation for understanding their functional importance during development. It has been observed that the overexpression of a 2-cell (2C) specific murine ERV in ES cells is sufficient to generate a 2C cellular phenotype (Macfarlan et al., 2012) This work suggests that TEs have the capacity to drive cell lineage specification. While differentially expressed TEs have been identified across different developmental stages of the preimplantation embryo (Goke et al., 2015), TE expression profiles have not been well characterized across different early stem cell lineages. ES and XEN stem cells have been reported to silence some TE expression, while TS cells are permissive for expression (Rowe et al., 2010; Golding et al., 2010). Based on the differences in their transcription regulatory networks, it is conceivable that different transposable elements classes are differently regulated in ES, TS, and XEN cells. Here we characterize the transposable element expression profiles of ES, TS, and XEN cells to test if specific families of transposable elements are associated with stem cell lineage specification.

## 2. METHODS

### 2.1. Cell Culture

Mouse ES-E14TG2a cells (ATCC CRL-1821) were maintained in 50% ESGRO-2i medium (MilliporeSigma) and 50% ES medium [DMEM (Gibco) with 15% fetal bovine serum (FBS) (Gibco), 100 μM nonessential amino acids (Gibco), 1 mM sodium pyruvate (Gibco), 55 μM 2-mercaptoethanol (Gibco), 2 mM L-glutamine (Gibco), 100 μg/mL penicillin, 100 μg/mL streptomycin, and 1000 μg/mL leukemia inhibitory factor (Gibco)] as previously described (Asanoma et al., 2011). Mouse TS cells were obtained from Dr. Janet Rossant (Hospital for Sick Children, Toronto, Canada) and maintained in FGF4/heparin supplemented TS medium [containing 30% TS basal medium (RPMI 1640 (Gibco) with 20% FBS, 1 mM sodium pyruvate, 100 μM 2-mercaptoethanol, 2 mM L-glutamine, 50 μg/mL penicillin, and 50 μg/mL streptomycin), 70% mouse embryonic fibroblast-conditioned medium, 25 ng/mL human recombinant FGF4 (Gibco), and 1 μg/mL heparin (Sigma-Aldrich)] as previously described (Tanaka et al., 1998).

### 2.2. RNA-Sequencing

Total RNA was extracted from the lysates of two wildtype ES cell replicates and five wildtype TS cell replicates with TRIzol (Invitrogen). 500 ng of total RNA was used for each RNA-seq library preparation. cDNA libraries were prepared using a TruSeq Stranded mRNA kit (Illumina) following the manufacturer’s instructions. mRNA was enriched from total RNA by oligo-dT magnetic beads, purified, and chemically fragmented. The first strand of cDNA was synthesized using random hexamer primers and Superscript II Reverse Transcriptase (Invitrogen). Second strand cDNA synthesis removed the RNA template and synthesized a replacement strand, incorporating dUTP in place of dTTP to generate double stranded (ds) cDNA. AMPure XP beads (Beckman Coulter) were used to purify the ds cDNA from the second strand reaction. The ds cDNA ends were blunted and poly(A) tails were added to the 3’ ends. After ligation of indexing adaptors (Illumina), the ds cDNA fragments were PCR amplified for 15 cycles. The cDNA libraries were sequenced on an Illumina HiSeq X platform (Novogene Corporation, Sacramento, CA).

### 2.3. Data Acquisition

Publicly available RNA-seq data of wild type ES (n = 3) and XEN (n = 7) cells from two studies (GSE61102, GSE106158 (Zhong et al., 2018)) were downloaded from the NCBI’s Gene Expression Omnibus (GEO) (Barrett et al., 2013). RNA-seq data collected from ES cells (n = 2) and TS cells (n = 5) for this study were also included with the downloaded data, leading to total sample sizes of n = 5 for ES data, n = 5 for TS data, and n = 7 for XEN data.

### 2.4. Transposable Element Analysis

Genomic coordinates of all repetitive elements were downloaded from the UCSC Genome Browser (Casper et al., 2018). Elements classified as “LTR” or “LINE” were kept while other elements were discarded because of the ability of LTRs and LINEs to autonomously replicate. Using the GenomicRanges (Lawrence et al., 2013), Biostrings, rtracklayer (Lawrence et al., 2009), and BSgenome.Mmusculus.UCSC.mm10 packages downloaded from Bioconductor, a FASTA file was generated with the sequences of all the filtered TEs which was later used to build a Kallisto index. Furthermore, the protein coding potentials of the filtered TEs were assessed with Coding Potential Calculator 2 (CPC2) (Kang et al., 2017). Sequences with a coding probability of 1 (i.e. 100%) were deemed protein-coding.

### 2.5. RNA-Seq Analysis

Using Kallisto (Bray et al., 2016), RNA-seq data were aligned with 5 bootstraps to a Kallisto index built from the mouse reference genome (mm10) downloaded from the UCSC Genome Browser. Transcript abundances from the Kallisto quantification were normalized as transcripts per million (TPM) using Sleuth (Pimentel et al., 2017). Each transcript was then assigned its gene name based on its RefSeq ID. To quantify TE abundance, RNA-seq data were aligned with 5 bootstraps to a Kallisto index built from the aforementioned filtered list of TEs using Kallisto. TE abundances from the Kallisto quantification were normalized and compared between each pair of groups (ES vs. TS, ES vs. XEN, and TS vs. XEN) with differential expression analysis using Sleuth. TEs with absolute beta value (analogous to absolute log transformed fold change) greater than 2 and FDR adjusted p-value (q-value) less than 0.05 were deemed differentially expressed. Additionally, the differential expression analysis results from each of the three comparisons were reanalyzed after being filtered to include only the aforementioned protein-coding TEs.

### 2.6. Statistical Analysis

Correlation between RNA-seq samples was determined with Pearson’s correlation coefficient. For comparisons among three means, one-way ANOVA was used followed by Tukey’s HSD post hoc test. The relationship between two categorical variables was tested using chi-square tests of independence. The α-level for all tests was at 0.05, meaning p < 0.05 was deemed significant. All statistical tests were performed in R.

## 3. RESULTS

### 3.1. Stem Cell Marker Expression Confirms RNA-Seq Sample Identity

Gene expression in each ES, TS, and XEN RNA-seq sample was quantified and analyzed. The validity of each sample’s lineage identity was assessed based on the sample’s expression of stem cell markers (*Foxd3, Nanog, Pou5f1*, and *Zfp42* for ES; *Cdx2, Elf5, Eomes*, and *Gata3* for TS; *Gata4, Gata6, Sox7*, and *Sox17* for XEN) (Lin et al., 2016; Ralston et al., 2010; Takahashi and Yamanaka, 2006). Based on a Pearson correlation matrix, there was high positive correlation among all XEN samples, among all TS samples, and among all ES samples (**Figure 1A**). Furthermore, the mean expression (TPM) of each gene’s most abundant transcript in all three cell types (**Figure 1B**) show the high expression of *Cdx2, Elf5, Eomes*, and *Gata3* in TS data relative to ES and XEN data, of *Foxd3, Nanog, Pou5f1*, and *Zfp42* in ES data relative to TS and XEN data, and of *Gata4, Gata6, Sox7*, and *Sox17* in XEN data relative to TS and ES data. These differences were confirmed to be significant (p < 0.05) through ANOVA and post hoc tests.

**Figure 1.**
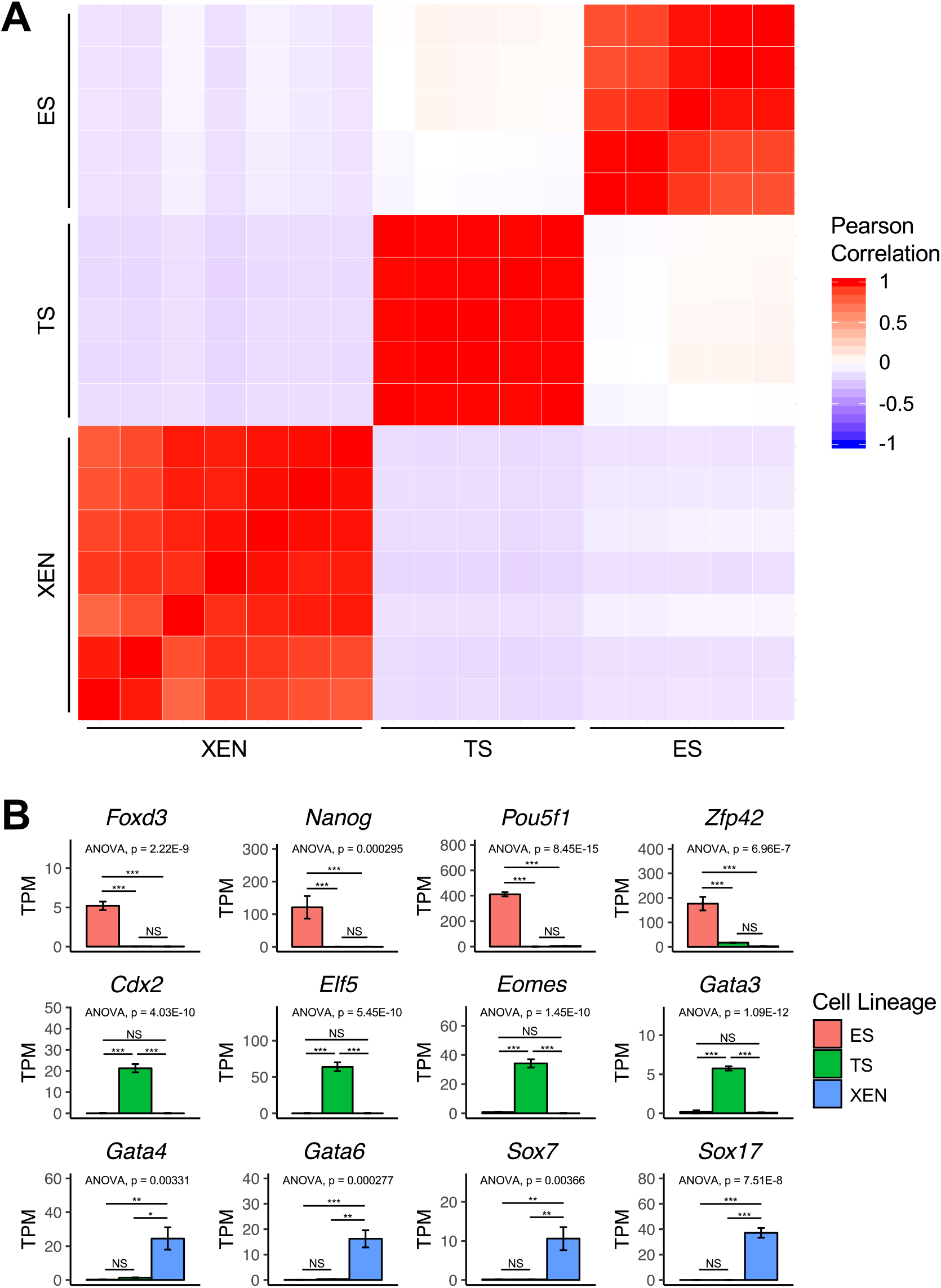
Stem Cell Marker Expression Confirms RNA-Seq Sample Identity. (A) A matrix shows the Pearson correlation of every RNA-seq sample with every other sample based on stem cell marker expression. (B) The mean expressions of the most abundant transcript of these markers (*Foxd3, Nanog, Pou5f1, Zfp42, Cdx2, Elf5, Eomes, Gata3, Gata4, Gata6, Sox7*, and *Sox17*) from the ES, TS, and XEN RNA-seq data are graphed. Additionally, ANOVA results for each gene’s expression among the cell types are shown on each graph. Error bars represent ± SEM, *** indicates p < 0.001, ** indicates p < 0.01, * indicates p < 0.05, and NS indicates no significant difference.

### 3.2. Differential Expression Analysis Reveals Distinct TE Expression Patterns in Early Stem Cell Lineages

TE expression in the ES, TS, and XEN RNA-seq data was quantified and compared through differential expression analysis in a pairwise manner: ES vs. TS, ES vs. XEN, and TS vs. XEN. From the differential expression analysis, volcano plots (**Figure 2A**) were generated showing the tendency for TEs to be upregulated in TS data in the ES vs. TS comparison, in neither ES nor XEN data in the ES vs. XEN comparison, and in TS data in the TS vs. XEN comparison. These tendencies are further illustrated in pie charts (**Figure 2B**) of all TEs from each analysis based on which cell line the TE is upregulated in. In the ES vs. TS analysis, 2.1% of TEs are upregulated in ES data compared to 12.8% in TS data. In the ES vs. XEN analysis, 7.0% of TEs are upregulated in ES data compared to 5.5% in XEN data. In the TS vs. XEN analysis, 15.1% of TEs are upregulated in TS data compared to 1.6% in XEN data. However, when the differentially expressed TEs in each analysis are sorted by TE family, there appears to be a tendency for the TEs of each family to be upregulated in a specific cell type (**Figures 2C and 2D**). This trend appeared from the ES vs. TS, ES vs. XEN, and TS vs. XEN analyses. Based on a chi-square test of independence of the differentially expressed TEs from the ES vs. TS analysis, there is a significant association between cell lineage and family of the differentially expressed TEs (χ^2^ = 1812.1, df = 377, p = 6.08 x 10^−186^), indicating families of TEs tend to be upregulated in a lineage-specific manner and confirming trends observed in **Figures 2C and 2D**. Similarly significant relationships between cell lineage and TE family were found in chi-square tests of the differentially expressed TEs from the ES vs. XEN analysis (χ^2^ = 367.80, df = 189, p = 1.34 x 10^−13^) and from the TS vs. XEN analysis (χ^2^ = 689.91, df = 366, p = 3.66 x 10^−22^).

**Figure 2.**
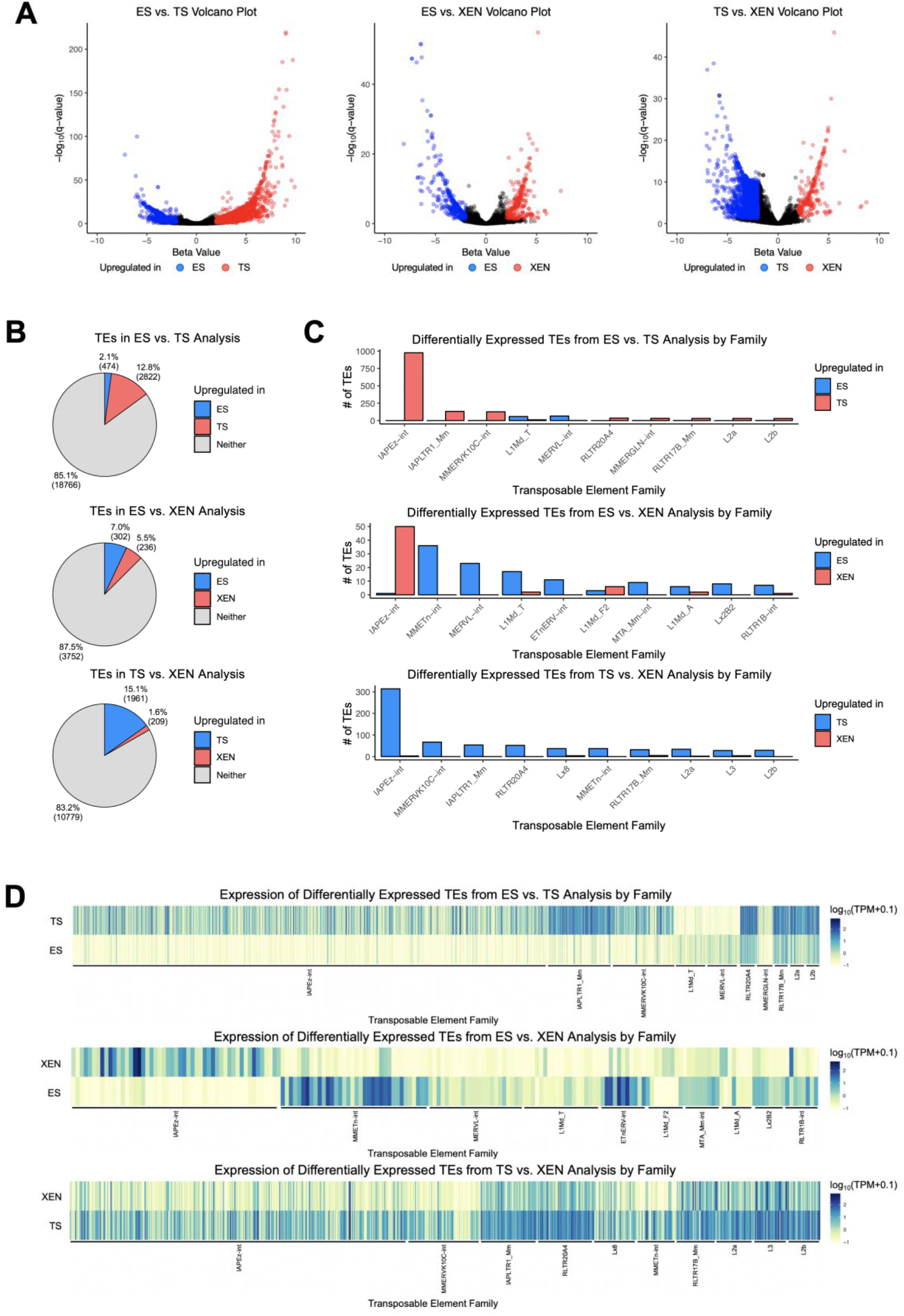
Differential Expression Analysis Reveals Distinct TE Expression Patterns in Early Stem Cell Lineages. (A) Volcano plots show the results of the ES vs. TS, ES vs. XEN, and TS vs. XEN differential expression analyses with every analyzed TE’s significance expressed as the negative of the log_10_ transformed q-value plotted against its beta value. Blue or red points represent differentially expressed TEs. (B) Pie charts show the proportions and frequencies of TEs upregulated in each cell type for each comparison. (C) From each analysis, the frequencies of differentially expressed TEs in either cell type are graphed by TE family. The 10 TE families with the greatest total number of differentially expressed TEs are shown for each analysis. (D) Heatmaps for each analysis show expression as measured by log_10_ transformed TPM with an added pseudocount of 0.1 of all TEs shown in (C) sorted and ordered by family in the same manner as (C).

### 3.3. TE Families with Protein-Coding Sequences Overlap with Potentially Lineage-Specific TE Families

The protein coding potential of all TEs used in the differential expression analysis was assessed using CPC2. After deeming TEs with 100% coding probability as protein-coding, it was revealed that the TE families with the highest number of integrations in the mouse genome (**Figure 3A**) vary from the families with the highest number of protein-coding integrations (**Figure 3B**). When comparing the results of this analysis with the results of the differential expression analysis, it was observed that several TE families which appeared to be lineage-specifically upregulated such as ETnERV, IAPEz, L1Md_A, and L1Md_T (**Figure 2C**) were also families containing the highest counts (**Figure 3B**) and highest proportions (**Figure 3C**) of protein-coding integrations. As such, it was predicted that differential expression analysis between the cell lineages based on only protein-coding TEs would show even more robust lineage-specific expression of TE families.

**Figure 3.**
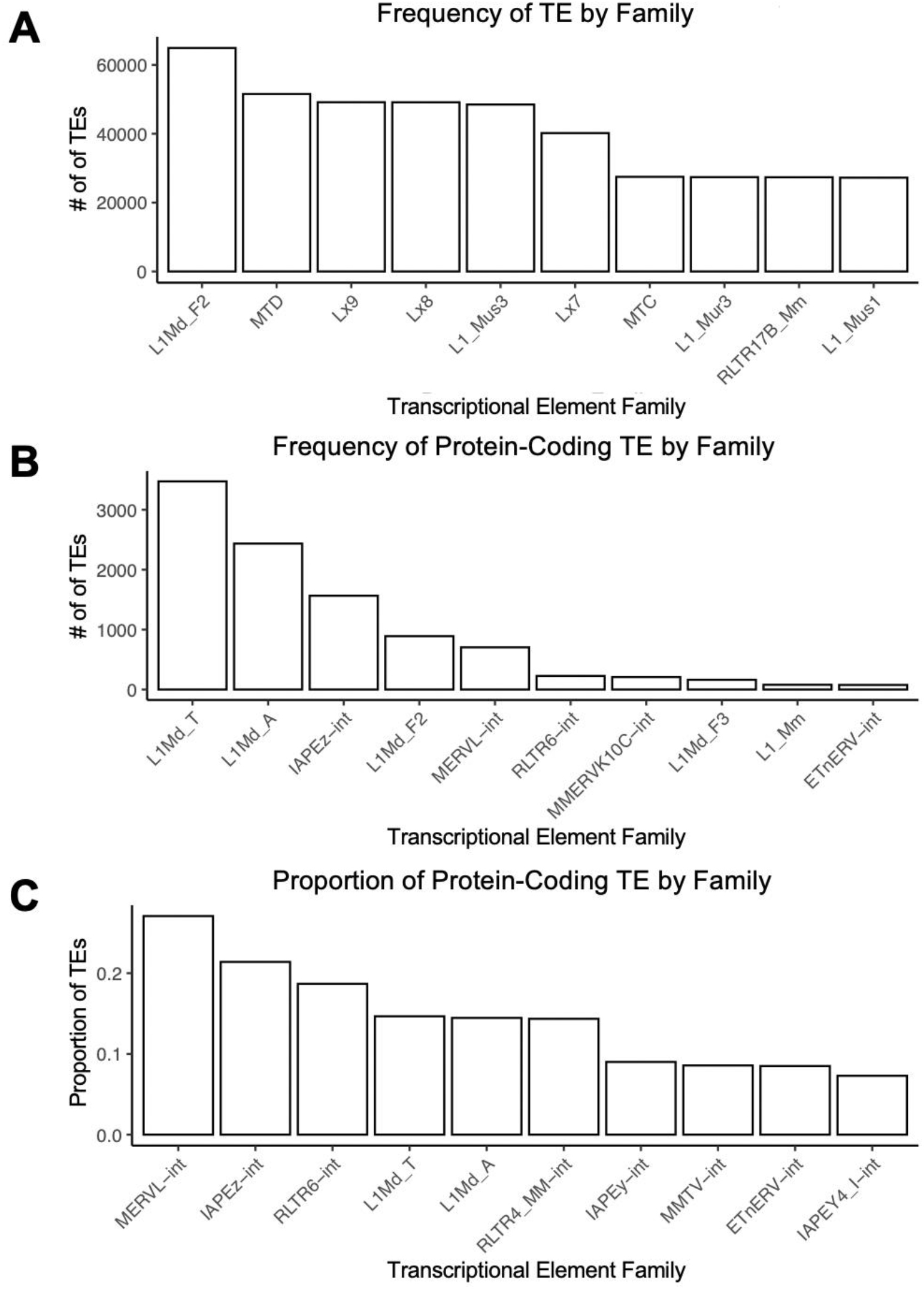
TE Families with Protein-Coding Sequences Overlap with Potentially Lineage-Specific TE Families. (A) The top 10 TE families with the highest numbers of integrations in the mouse genome. (B) The top 10 TE families with the highest numbers of protein-coding integrations in the mouse genome. (C) The top 10 TE families with the highest proportions of protein-coding TE integrations versus all TE integrations.

### 3.4. Differential Expression Analysis of Protein-Coding TEs Reveals Lineage-Specific Protein-Coding TE Families

The results of the previous differential expression analyses (**Figure 2**) were reanalyzed by filtering each set of results to include only TEs with the potential to encode protein. Using these filtered results, volcano plots (**Figure 4A**) were generated showing the tendency for protein-coding TEs to be upregulated in TS data in the ES vs. TS comparison, in neither ES nor XEN data in the ES vs. XEN comparison, and in TS data in the TS vs. XEN comparison. These tendencies of protein-coding TE expression based on cell type appear to be more pronounced than the patterns observed from all TEs (**Figure 2A**). The more distinct expression patterns of protein-coding TEs are illustrated in pie charts (**Figure 4B**) of all protein-coding TEs from each analysis based on which cell line the protein-coding TE is upregulated in. In the ES vs. TS analysis, 8.9% of TEs are upregulated in ES data compared to 58.8% in TS data. In the ES vs. XEN analysis, 16.4% of TEs are upregulated in ES data compared to 12.9% in XEN data. In the TS vs. XEN analysis, 26.5% of TEs are upregulated in TS data compared to 0.8% in XEN data. These data also serve to show the greater tendency for protein-coding TEs to be differentially expressed as compared to TEs in general. When the differentially expressed protein-coding TEs in each analysis are sorted by TE family, there appears to be tendency for the protein-coding TEs of each family to be upregulated in a specific cell type (**Figures 4C and 4D**). This trend appeared from the ES vs. TS, ES vs. XEN, and TS vs. XEN analyses. Based on a chi-square test of independence of the differentially expressed protein-coding TEs from the ES vs. TS analysis, there is a significant association between cell lineage and family of the differentially expressed protein-coding TEs (χ^2^ = 820.97, df = 14, p = 3.61 x 10^−166^), indicating families of protein-coding TEs tend to be upregulated in a lineage-specific manner. From this test, it was also seen that ETnERV, L1Md_A, L1Md_T, L1_Mus1, MERVL_2A, MERVL, MMETn, and RLTR4_MM sequences upregulated in ES cells and IAPEz and MMERVK10C sequences upregulated in TS cells are observed significantly more than expected (absolute value of standardized residual > 2). Similarly significant relationships between cell lineage and TE family were found in chi-square tests of the differentially expressed TEs from the ES vs. XEN analysis (χ^2^ = 93.01, df = 13, p = 3.71 x 10^−14^) and from the TS vs. XEN analysis (χ^2^ = 232.76, df =16, p = 1.75 x 10^−40^). From the ES vs. XEN test, L1Md_T, MERVL, MMETn, and RLTR4_MM sequences upregulated in ES cells and IAPEY4_I and IAPEz sequences upregulated in XEN cells were observed significantly more than expected. From the TS vs. XEN test, IAPEz sequences upregulated in TS cells and IAPEY4_I and L1Md_F2 sequences upregulated in XEN were observed significantly more than expected. Considering that TE families with a tendency to be upregulated in one lineage relative to the other two are likely to be lineage-specifically expressed, IAPEz expression is likely specific to TS cells, IAPEY4_I expression is likely specific to XEN cells, and L1Md_T, MERVL, MMETn, and RLTR4_MM expression is likely specific to ES cells.

**Figure 4.**
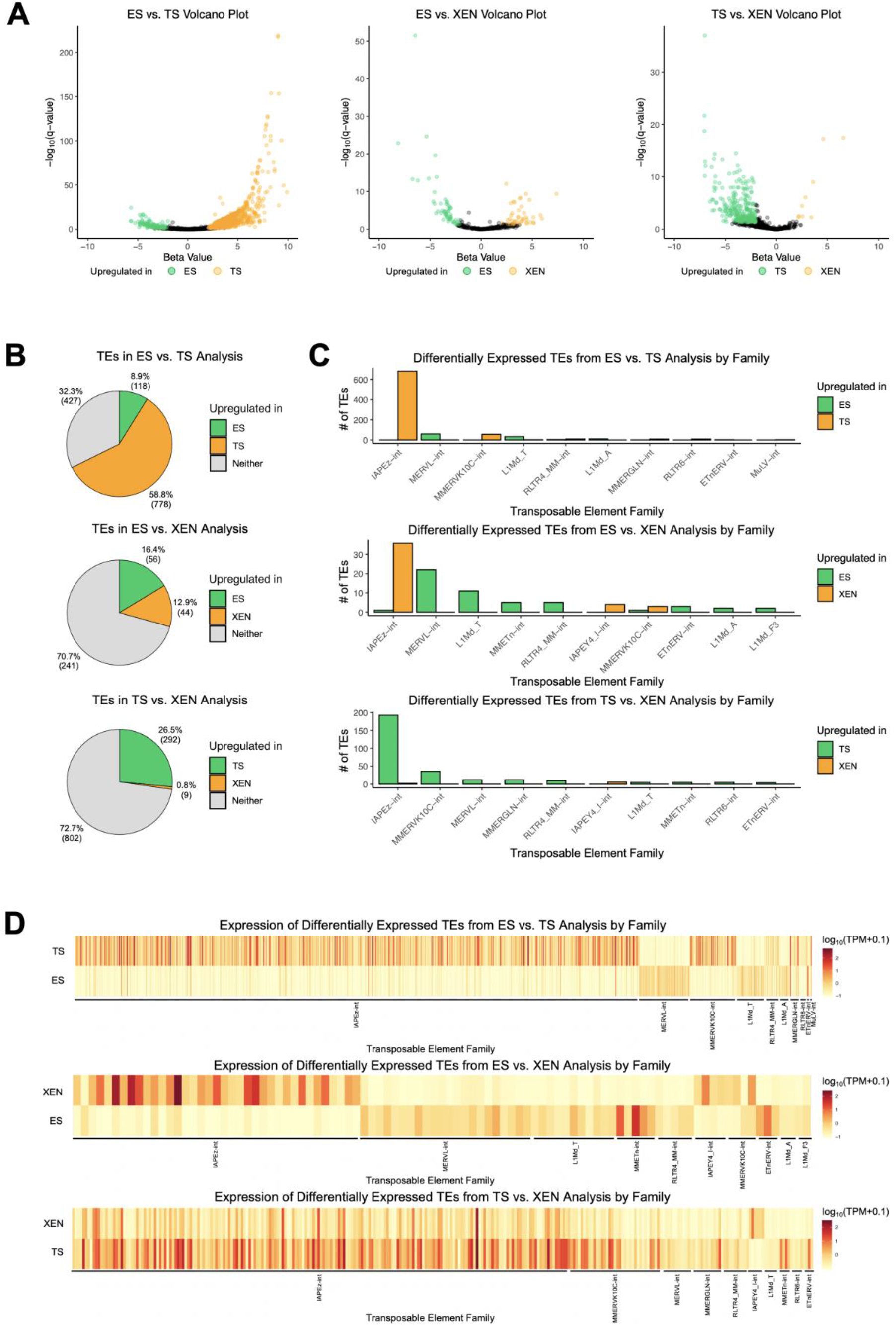
Differential Expression Analysis of Protein-Coding TEs Reveals Lineage-Specific Protein-Coding TE Families. (A) Volcano plots show the results of the ES vs. TS, ES vs. XEN, and TS vs. XEN differential expression analyses with every analyzed protein-coding TE’s significance expressed as the negative of the log_10_ transformed q-value plotted against its beta value. Blue or red points represent differentially expressed protein-coding TEs. (B) Pie charts show the proportions and frequencies of protein-coding TEs upregulated in each cell type for each comparison. (C) From each analysis, the frequencies of differentially expressed protein-coding TEs in either cell type are graphed by TE family. The 10 TE families with the greatest total number of differentially expressed protein-coding TEs are shown for each analysis. (D) Heatmaps for each analysis show expression as measured by log_10_ transformed TPM with an added pseudocount of 0.1 of all protein-coding TEs shown in (C) sorted and ordered by family in the same manner as (C).

## 4. DISCUSSION

In this study, the expression of TEs in the transcriptomes of mouse ES, TS, and XEN cells was analyzed and compared to better understand what function TEs may serve in early embryonic development and stem cell differentiation. Using both RNA-seq data generated for this study and obtained from previous studies, the abundance of LTR and LINE sequences was quantified in each sample and compared pairwise between the cell lineages. Differential expression analysis between ES and TS data, ES and XEN data, and TS and XEN data revealed TE expression is generally higher in TS data relative to both ES and XEN data but that TE families within these analyses are lineage-specifically upregulated (**Figure 2**). We also observed that potential protein coding transposable elements are cell type restricted (**Figure 4**).

One possible explanation for the observed lineage-specific expression of many TE families could be the upregulation of these families by lineage-specific transcription factors (TFs), e.g. NANOG and POU5F for ES cells, CDX2 and ELF5 for TS cells, GATA4 and GATA6 for XEN cells (Lin et al., 2016; Ralston et al., 2010; Takahashi and Yamanaka, 2006). It has been reported that TFs may be binding to a promoter or enhancer region either within or near particular TE loci, driving transcription of those TEs (Goke et al., 2015). Using the results of this study, chromatin immunoprecipitation sequencing (ChIP-seq) data of such lineage-specific TFs could be analyzed for regions upstream of, within, and downstream of the loci of differentially expressed TEs, similar to previous studies (Chuong et al., 2013), to study if lineage-specific TE expression is driven by these TFs. Another potential implication of the identified lineage-specifically expressed TEs is their possible functional role in stem cell differentiation. TEs have been shown to have a regulatory role in ES cell maintenance (Fort et al., 2014), but TE-encoded proteins have also been shown to be important for such maintenance (Macfarlan et al., 2012).

TEs from the intracisternal A-type particle (IAP) family, a type of ERV, are increasingly expressed from the zygote up to the blastocyst stage of the preimplantation mouse embryo (Svoboda et al., 2004), indicating that IAPs may be important for stem cell lineage determination, development, and maintenance. In this study, it was observed that many IAPEz TEs are specifically expressed in TS cells while many IAPEY4_I TEs are specifically expressed in XEN cells. In contrast, there is a lack of any IAP expression in ES cells. A possible explanation for this observation may be that the silencing of IAPs is involved in ES lineage determination or cellular function. It has been demonstrated that deletion of KAP1 in ES cells leads to an increase of ERVs expression and particularly IAPs (Rowe et al., 2010), indicating that IAPs are normally silenced in ES cells as was supported here. Another possibility may be that the expression of IAP sequences and the possible proteins they encode play a role in TS and/or XEN cell development, similar to how env-derived syncitin proteins from ERVs are necessary for mammalian placenta formation (Blaise et al., 2005). This is supported by the fact that IAPs have been shown to be both mobile (Dewannieux et al., 2004) and capable of coding for full retroviral proteins (Ribet et al., 2008a) in mouse cells.

While IAPEz were shown to be expressed in TS and IAPEY4_I in XEN cells, various other TE families were shown to be specifically expressed in ES cells including L1Md_T, MERVL, MMETn, and RLTR4_MM. L1Md_T is a subfamily of LINE-1, a family containing sequences which previous studies have shown to be expressed in ES cells (Garcia-Perez et al., 2007). There is a possibility that LINE-1 sequences may have a role in ES lineage determination, supported by previous findings that LINE-1 sequences have two open reading frames (ORFs), with ORF1 encoding proteins that serve as nucleic acid chaperones (Kolosha and Martin, 1997; Martin and Bushman, 2001) and ORF2 encoding reverse transcriptase (Feng et al., 1996; Mathias et al., 1991). Another ES-specific family is MERVL, also known as MuERV-L. The ERVL family exists in all placental mammals and was thus present in the mammalian common ancestor more than 70 million years ago (mya) (Benit et al., 1999). Despite its age, the MERVL family has maintained some active elements in the mouse, as proven by recent amplifications (Benit et al., 1999; Costas, 2003). It has also been shown that MERVL sequences encode *gag* and *pol* genes but no *env* gene and produce virus-like particles in the early mouse embryo (Ribet et al., 2008b), consistent with this study and others showing high levels of expression during early embryonic development (Svoboda et al., 2004). This study also shows that MERVL sequences are specifically expressed in the ES lineage, indicating the possibility that MERVL-encoded proteins may be involved in ES lineage determination which is further supported by the ability for MERVL expression to increase cell potency (Macfarlan et al., 2012). While there has been little previous study of the MMETn (*Mus musculus* early transposon) and RLTR4_MM families, protein-coding sequences from these families were also observed to be specifically expressed in ES cells, indicating that they may be involved in ES differentiation.

TEs are species-specific, meaning the implications of this study most directly apply to the role of TE expression in mouse embryonic development (Brind’Amour et al., 2018; Robbez-Masson and Rowe, 2015). However, this study has implications concerning the potential importance of TE in the embryonic development of other species. For humans, this can be confirmed by applying the analyses in this study to human early stem cell transcriptomic data as it becomes increasingly prevalent. Nevertheless, this study further enforces the theory that TEs play a role in early embryonic development and also establishes that TEs may contribute to stem cell differentiation in the early embryo.

## ACKNOWLEDGEMENTS

This study was supported by the funding from NIH/NICHD grant HD079363.

## REFERENCES

Asanoma, K., Rumi, M.A.K., Kent, L.N., Chakraborty, D., Renaud, S.J., Wake, N., Lee, D.-S., Kubota, K., and Soares, M.J. (2011). FGF4-dependent stem cells derived from rat blastocysts differentiate along the trophoblast lineage. Developmental biology 351, 110–119.

Baker, C.L., and Pera, M.F. (2018). Capturing Totipotent Stem Cells. Cell stem cell 22, 25–34.

Barrett, T., Wilhite, S.E., Ledoux, P., Evangelista, C., Kim, I.F., Tomashevsky, M., Marshall, K.A., Phillippy, K.H., Sherman, P.M., Holko, M., et al. (2013). NCBI GEO: archive for functional genomics data sets--update. Nucleic acids research 41, D991–995.

Benit, L., Lallemand, J.B., Casella, J.F., Philippe, H., and Heidmann, T. (1999). ERV-L elements: a family of endogenous retrovirus-like elements active throughout the evolution of mammals. Journal of virology 73, 3301–3308.

Blaise, S., de Parseval, N., and Heidmann, T. (2005). Functional characterization of two newly identified Human Endogenous Retrovirus coding envelope genes. Retrovirology 2, 19.

Bray, N.L., Pimentel, H., Melsted, P., and Pachter, L. (2016). Near-optimal probabilistic RNA-seq quantification. Nature biotechnology 34, 525–527.

Brind’Amour, J., Kobayashi, H., Richard Albert, J., Shirane, K., Sakashita, A., Kamio, A., Bogutz, A., Koike, T., Karimi, M.M., Lefebvre, L., et al. (2018). LTR retrotransposons transcribed in oocytes drive species-specific and heritable changes in DNA methylation. Nat Commun 9, 3331–3331.

Casper, J., Zweig, A.S., Villarreal, C., Tyner, C., Speir, M.L., Rosenbloom, K.R., Raney, B.J., Lee, C.M., Lee, B.T., Karolchik, D., et al. (2018). The UCSC Genome Browser database: 2018 update. Nucleic acids research 46, D762–d769.

Chuong, E.B., Rumi, M.A., Soares, M.J., and Baker, J.C. (2013). Endogenous retroviruses function as species-specific enhancer elements in the placenta. Nature genetics 45, 325–329.

Costas, J. (2003). Molecular characterization of the recent intragenomic spread of the murine endogenous retrovirus MuERV-L. J Mol Evol 56, 181–186.

Dewannieux, M., Dupressoir, A., Harper, F., Pierron, G., and Heidmann, T. (2004). Identification of autonomous IAP LTR retrotransposons mobile in mammalian cells. Nature genetics 36, 534–539.

Feng, Q., Moran, J.V., Kazazian, H.H., Jr., and Boeke, J.D. (1996). Human L1 retrotransposon encodes a conserved endonuclease required for retrotransposition. Cell 87, 905–916.

Feschotte, C., and Gilbert, C. (2012). Endogenous viruses: insights into viral evolution and impact on host biology. Nat Rev Genet 13, 283–296.

Finnegan, D.J. (2012). Retrotransposons. Current Biology 22, R432–R437.

Fort, A., Hashimoto, K., Yamada, D., Salimullah, M., Keya, C.A., Saxena, A., Bonetti, A., Voineagu, I., Bertin, N., Kratz, A., et al. (2014). Deep transcriptome profiling of mammalian stem cells supports a regulatory role for retrotransposons in pluripotency maintenance. Nature genetics 46, 558–566.

Garcia-Perez, J.L., Marchetto, M.C., Muotri, A.R., Coufal, N.G., Gage, F.H., O’Shea, K.S., and Moran, J.V. (2007). LINE-1 retrotransposition in human embryonic stem cells. Human molecular genetics 16, 1569–1577.

Gifford, R., and Tristem, M. (2003). The evolution, distribution and diversity of endogenous retroviruses. Virus Genes 26, 291–315.

Goke, J., Lu, X., Chan, Y.S., Ng, H.H., Ly, L.H., Sachs, F., and Szczerbinska, I. (2015). Dynamic transcription of distinct classes of endogenous retroviral elements marks specific populations of early human embryonic cells. Cell stem cell 16, 135–141.

Golding, M.C., Zhang, L., and Mann, M.R. (2010). Multiple epigenetic modifiers induce aggressive viral extinction in extraembryonic endoderm stem cells. Cell stem cell 6, 457–467.

Jacques, P.E., Jeyakani, J., and Bourque, G. (2013). The majority of primate-specific regulatory sequences are derived from transposable elements. PLoS genetics 9, e1003504.

Jern, P., and Coffin, J.M. (2008). Host-retrovirus arms race: trimming the budget. Cell Host Microbe 4, 196–197.

Joly-Lopez, Z., and Bureau, T.E. (2018). Exaptation of transposable element coding sequences. Current opinion in genetics & development 49, 34–42.

Kang, Y.J., Yang, D.C., Kong, L., Hou, M., Meng, Y.Q., Wei, L., and Gao, G. (2017). CPC2: a fast and accurate coding potential calculator based on sequence intrinsic features. Nucleic acids research 45, W12–w16.

Kolosha, V.O., and Martin, S.L. (1997). In vitro properties of the first ORF protein from mouse LINE-1 support its role in ribonucleoprotein particle formation during retrotransposition. Proceedings of the National Academy of Sciences of the United States of America 94, 10155–10160.

Lander, E.S., Linton, L.M., Birren, B., Nusbaum, C., Zody, M.C., Baldwin, J., Devon, K., Dewar, K., Doyle, M., FitzHugh, W., et al. (2001). Initial sequencing and analysis of the human genome. Nature 409, 860–921.

Lawrence, M., Gentleman, R., and Carey, V. (2009). rtracklayer: an R package for interfacing with genome browsers. Bioinformatics 25, 1841–1842.

Lawrence, M., Huber, W., Pages, H., Aboyoun, P., Carlson, M., Gentleman, R., Morgan, M.T., and Carey, V.J. (2013). Software for computing and annotating genomic ranges. PLoS computational biology 9, e1003118.

Lin, J., Khan, M., Zapiec, B., and Mombaerts, P. (2016). Efficient derivation of extraembryonic endoderm stem cell lines from mouse postimplantation embryos. Scientific reports 6, 39457.

Macfarlan, T.S., Gifford, W.D., Driscoll, S., Lettieri, K., Rowe, H.M., Bonanomi, D., Firth, A., Singer, O., Trono, D., and Pfaff, S.L. (2012). Embryonic stem cell potency fluctuates with endogenous retrovirus activity. Nature 487, 57–63.

Martin, S.L., and Bushman, F.D. (2001). Nucleic acid chaperone activity of the ORF1 protein from the mouse LINE-1 retrotransposon. Molecular and cellular biology 21, 467–475.

Mathias, S.L., Scott, A.F., Kazazian, H.H., Jr., Boeke, J.D., and Gabriel, A. (1991). Reverse transcriptase encoded by a human transposable element. Science (New York, NY) 254, 1808–1810.

Meyer, T.J., Rosenkrantz, J.L., Carbone, L., and Chavez, S.L. (2017). Endogenous Retroviruses: With Us and against Us. Frontiers in Chemistry 5.

Naville, M., Warren, I.A., Haftek-Terreau, Z., Chalopin, D., Brunet, F., Levin, P., Galiana, D., and Volff, J.N. (2016). Not so bad after all: retroviruses and long terminal repeat retrotransposons as a source of new genes in vertebrates. Clinical microbiology and infection: the official publication of the European Society of Clinical Microbiology and Infectious Diseases 22, 312–323.

Pimentel, H., Bray, N.L., Puente, S., Melsted, P., and Pachter, L. (2017). Differential analysis of RNA-seq incorporating quantification uncertainty. Nature methods 14, 687–690.

Ralston, A., Cox, B.J., Nishioka, N., Sasaki, H., Chea, E., Rugg-Gunn, P., Guo, G., Robson, P., Draper, J.S., and Rossant, J. (2010). Gata3 regulates trophoblast development downstream of Tead4 and in parallel to Cdx2. Development (Cambridge, England) 137, 395–403.

Ribet, D., Harper, F., Dupressoir, A., Dewannieux, M., Pierron, G., and Heidmann, T. (2008a). An infectious progenitor for the murine IAP retrotransposon: emergence of an intracellular genetic parasite from an ancient retrovirus. Genome research 18, 597–609.

Ribet, D., Louvet-Vallee, S., Harper, F., de Parseval, N., Dewannieux, M., Heidmann, O., Pierron, G., Maro, B., and Heidmann, T. (2008b). Murine endogenous retrovirus MuERV-L is the progenitor of the “orphan” epsilon viruslike particles of the early mouse embryo. Journal of virology 82, 1622–1625.

Robbez-Masson, L., and Rowe, H.M. (2015). Retrotransposons shape species-specific embryonic stem cell gene expression. Retrovirology 12, 45–45.

Rowe, H.M., Jakobsson, J., Mesnard, D., Rougemont, J., Reynard, S., Aktas, T., Maillard, P.V., Layard-Liesching, H., Verp, S., Marquis, J., et al. (2010). KAP1 controls endogenous retroviruses in embryonic stem cells. Nature 463, 237–240.

Stoye, J.P. (2012). Studies of endogenous retroviruses reveal a continuing evolutionary saga. Nature reviews Microbiology 10, 395–406.

Svoboda, P., Stein, P., Anger, M., Bernstein, E., Hannon, G.J., and Schultz, R.M. (2004). RNAi and expression of retrotransposons MuERV-L and IAP in preimplantation mouse embryos. Developmental biology 269, 276–285.

Takahashi, K., and Yamanaka, S. (2006). Induction of pluripotent stem cells from mouse embryonic and adult fibroblast cultures by defined factors. Cell 126, 663–676.

Tanaka, S., Kunath, T., Hadjantonakis, A.-K., Nagy, A., and Rossant, J. (1998). Promotion of Trophoblast Stem Cell Proliferation by FGF4. Science (New York, NY) 282, 2072.

Wamaitha, S.E., and Niakan, K.K. (2018). Human Pre-gastrulation Development. Current topics in developmental biology 128, 295–338.

Zhong, Y., Choi, T., Kim, M., Jung, K.H., Chai, Y.G., and Binas, B. (2018). Isolation of primitive mouse extraembryonic endoderm (pXEN) stem cell lines. Stem cell research 30, 100–112.

